# Electrophysiological characterization of medial preoptic neurons in lactating rats and its modulation by hypocretin-1

**DOI:** 10.1101/2022.06.21.497055

**Authors:** Mayda Rivas, Diego Serantes, Claudia Pascovich, Florencia Peña, Annabel Ferreira, Pablo Torterolo, Luciana Benedetto

## Abstract

The medial preoptic area (mPOA) undergoes through neuroanatomical changes across the postpartum period, during which its neurons play a critical role in the regulation of maternal behavior. In addition, this area is also crucial for sleep-wake regulation. We have previously shown that hypocretins (HCRT) within the mPOA facilitate active maternal behaviors in postpartum rats, while the blockade of endogenous HCRT in this area promotes nursing and sleep. To explore the mechanisms behind these HCRT actions, we aimed to evaluate the effects of juxta-cellular HCRT-1 administration on mPOA neurons in urethane-anesthetized postpartum and virgin female rats. We recorded mPOA single units and the electroencephalogram (EEG) and applied HCRT-1 juxta-cellular by pressure pulses. Our main results show that the electrophysiological characteristics of the mPOA neurons and their relationship with the EEG of postpartum rats did not differ from virgin rats. Additionally, neurons that respond to HCRT-1 had a slower firing rate than those that did not. In addition, administration of HCRT increased the activity in one group of neurons while decreasing it in another, both in postpartum and virgin rats. The mechanisms by which HCRT modulate functions controlled by the mPOA involve different cell populations.

## Introduction

The medial preoptic area (mPOA) is a crucial hypothalamic region that controls sleep (Benedetto et al., 2021; Vanini and Torterolo, 2021), thermoregulation (Kumar, 2004), and maternal care (Numan, 1974; Numan, 2006). During the transition to motherhood and postpartum period, the mPOA undergoes through anatomical and physiological changes that allow the female to adapt to the maternal care of the pups. In this sense, it has been described modifications in the complexity of dendritic spines in maternal mPOA neurons (Shams et al., 2012; Parent et al., 2017), in the expression of hormone receptors such as estradiol, progesterone, prolactin, and oxytocin (Insel, 1990; Wagner and Morrell, 1996; Numan et al., 1999; Mann and Bridges, 2001), as well as in the density of neurotransmitters’ receptors, including dopamine, vasopressin, neurotensin, amylin, and GABA (Gammie, 2005; Bosch and Neumann, 2008; Dobolyi, 2009; Akbari et al., 2013; Driessen et al., 2014). In addition, there are changes in the neurotransmitter phenotype of mPOA neurons, such as the expression of melanin-concentrating hormone (MCH) in GABAergic neurons, that only occur during the postpartum period (Rondini et al., 2010). However, it is unknown if these postpartum adaptations lead to electrophysiological changes in mPOA neurons.

mPOA neurons are critical for the normal development of maternal behaviors (Fleming and Walsh, 1994; Lonstein et al., 1998). In this sense, mPOA neurons increase their c-Fos expression (a marker for neuronal activity) when the mother displays active maternal behaviors (Fleming and Walsh, 1994; Lonstein et al., 1998) and remains high for two days after contact with the litter (Stack and Numan, 2000). Most of these active mPOA neurons are GABAergic (Tsuneoka et al., 2013), and recent evidence shows that chemogenetic activation of GABAergic preoptic neurons increases maternal behavior (Dimen et al., 2021). However, the electrophysiological characteristics of mPOA neurons during the postpartum period are mostly unknown. Only Fang et al. (2018) have studied the activity of mPOA neurons in mice at different reproductive states. These authors described that the neuronal firing rate decreases in lactating mothers compared to virgin and post-lactation animals. Given this precedent, together with the neuroanatomic-functional changes described above, we hypothesize that the activity of the mPOA neurons in lactating and virgin female rats differ in several electrophysiological parameters.

Sleep-active and wake-active neurons are intermingled within the mPOA of male mice and rats (Szymusiak et al., 1998; Takahashi et al., 2009). While most sleep-active neurons are GABAergic (Gong et al., 2004), Vanini and collaborators recently described a group of glutamatergic neurons in the ventral half of the mPOA, in which chemogenetic activation increases wakefulness and decreases both NREM and REM sleep (Vanini et al., 2020; Mondino et al., 2021). However, there are no studies regarding mPOA neuronal activity in the postpartum period during sleep.

The hypocretinergic (HCRTergic) system contributes to the maintenance of wakefulness and regulates several motivating activities, including maternal behavior (Torterolo et al., 2003; Harris et al., 2005; D’Anna and Gammie, 2006; Muschamp et al., 2007; Boutrel et al., 2010; McGregor et al., 2011; Torterolo et al., 2011; Rivas et al., 2016; Rivas et al., 2021). Hypocretin-1 (HCRT-1) and HCRT-2 (also called Orexins A and B, respectively) are neuropeptides synthesized by neurons located in the postero-lateral hypothalamus, which project to many brain areas, including the mPOA (Peyron et al., 1998; Trivedi et al., 1998). Interestingly, Fos immunoreactivity of HCRTergic neurons as well as the expression of pre-pro-HCRT mRNA and HCRT receptor-1 mRNA, increase during the postpartum period in mice and rats (Wang et al., 2003; Espana et al., 2004).

We have recently shown that HCRT-1 within the mPOA facilitates some active maternal behaviors in postpartum rats, and the blockade of endogenous HCRT in the mPOA promotes nursing and sleep (Rivas et al., 2016; Rivas et al., 2021). However, to our knowledge, the effect of HCRT on mPOA unit activity has not been previously reported.

In the present report, as a first step to characterize the *in vivo* electrophysiology of the mPOA neurons in postpartum and virgin rats, we utilized urethane anesthesia as a proxy for the sleep-wake cycle. Under this anesthesia, there are spontaneous and rhythmic alternations between two different electroencephalographic patterns, a slow-wave state that resembles NREM sleep, and an active state with features of both REM sleep and wakefulness (Clement et al., 2008; Pagliardini et al., 2013; Mondino et al., 2022). Mondino et al. (2022) named these urethane-induced states NREMure and REMure, respectively. In addition, we also evaluated the effect on neuronal activity of juxta-cellular administration of HCRT-1 in both lactating and virgin rats.

## Materials and Methods

### Animals and housing

We utilized fifteen diestrus virgin females and twenty-two primiparous lactating Wistar female rats weighing 230 to 350 g (days six to eight postpartum). Animals were housed in a temperature-controlled (22 ± 1 °C) room, under a 12-h light/dark cycle (lights on at 6:00 a.m.), with *ad libitum* access to food and water. The experimental procedures were approved by the Institutional Animal Care Committee (protocol nº 070153-000550-18).

### Stereotaxic surgery

The recordings were made in rats anesthetized with urethane (1.2 g/kg, i.p.), maintaining body temperature at 37.0-37.5 °C using a heating pad. We positioned the head of the rat in a stereotaxic frame, made a scalp incision, and exposed the skull. According to Paxinos and Watson (2005) coordinates (AP -0.5 mm, L 0.5 mm, H 8.0 – 9.3; from Bregma), we drilled a small hole in the skull to descend a recording electrode into the mPOA. Additional holes were performed to place three stainless-steel screw electrodes for electroencephalogram (EEG) recordings in the following locations: frontal cortex (AP = +3.5, ML = 2.0), occipital cortex (AP = -6.5, ML = 2.0) and cerebellum (AP = -11.0, ML = 0.0; as a reference electrode).

### Unit and EEG recordings

We carried out unit recordings using standard procedures with an Ag-Cl electrode inside a glass micropipette of 5-20 MΩ, filled with 2M NaCl; we also handmade fabricated double micropipettes for simultaneous recording and juxta-cellular administration of HCRT (Torterolo et al., 1998; Torterolo et al., 2002; Devera et al., 2015; Pascovich et al., 2020). An Ag-Cl electrode under the neck skin was used as reference for unit recordings. The HCRT-filled micropipette tip had a diameter of 5-20 mm and was separated from the recording electrode tip by 50-150 mm. Neuronal signals were amplified x 1000 by an AC-coupled amplifier (Dagan 2400A), filtered between 300 Hz - 10 kHz, and digitized at 20 kHz. Single unit activity in the mPOA was acquired and processed using Spike2 software (Cambridge Electronic Design, UK). Baseline firing of mPOA units was recorded for at least 300 s. Afterward, juxta-cellular HCRT-1 (100 μM, BACHEM, Bubendorf, Switzerland) diluted in sterile saline 0.9 % (or its vehicle) was applied by pressure pulses of 20 PSI for 200 to 300 ms (Devera et al., 2015; Pascovich et al., 2020). EEG signals were amplified (x1000), filtered (0.1–500 Hz), sampled (1024 Hz, 16 bits), acquired, and processed using the Spike2 software.

### Histological verification of unit recording sites

At the end of the experiment, the animal was euthanized with an overdose of urethane and perfused with 4% paraformaldehyde. The brain was removed and cut in coronal sections (150 μm) using a vibratome. Brain sections were observed under an optical microscope to localize micropipettes traces according to the neuroanatomical atlas of Paxinos and Watson (2005).

### Electrophysiological recording analysis

We sorted the single units according to amplitude and waveform criteria. We examined the lack of spikes during the refractory period (< 2 ms), confirming the absence of contamination by other units. Then, we analyzed the average waveforms of the action potentials (AP). The APs were mostly biphasic; the AP duration was considered between the beginning of the first phase and the end of second phase, despite a third phase was observed in some cases (Hajos et al., 1995; Devera et al., 2015).

The basal pattern of discharge was analyzed offline by mean and instantaneous frequency, their respective coefficients of variation (SD/mean), interval histogram (IH), and autocorrelation histogram (AH). In both histograms, bins of 1 ms were used. These parameters were calculated for each recorded unit from the total number of spikes occurring in the first 300 s of stable activity. In neurons whose activity varied according to EEG state, the basal rate was calculated during NREMure (EEG-slow waves state).

Recorded neurons were also categorized in different groups according to:

1. Reproductive state: lactating or virgin.
2. Firing pattern. Taking into account the IH, AH and the raw recording, four groups of neurons were defined: arrhythmic (neuronal discharge without a rhythm in the AH); rhythmic (rhythmic neuronal discharge profile in the AH); long-duration burst (rhythmic neuronal firing trains separated by intervals longer than 10 sec, with highly stereotyped firing seen in the raw recording); predominant interval (neurons with a main interval observable in both IH and AH; these neurons showed burst discharge; Figure 2).
3. EEG relation. The neurons were classified as those that increase their activity during NREMure (NREMure-ON), REMure (REMure-ON), or their activity was unrelated to the EEG state.
4. HCRT-1 effect. Neurons were categorized according to their response to HCRT-1 administration in increased, decreased, or no-change.

### Statistical analyses

For the statistical analyses we used SPSS 21 (IBM) and described the data as the mean and standard error of the mean (SEM). We employed the Generalized Linear Mixed Models (GLMM) to test the effects of predefined variables on electrophysiological parameters from the single-cell recordings, with gamma distribution and log as link functions. For categorical response variables, binomial distribution was used in presence or absence as in burst firing, and multinomial distribution for several categories as in firing patterns of neurons (Agresti, 2003). We used random effects for animals since we recorded several cells in the same rat, but fixed effects for reproductive condition, EEG relationship, and HCRT results. Bonferroni *post hoc* test (sequential adjusted) was used for pairwise comparisons. Information criteria for the selection of models were based on the –2 log pseudo-likelihood in SPSS.

We used the Mann–Whitney test to compare the firing rate of each neuron during NREMure vs. REMure (windows of 1 to 5 minutes depending on the firing rate of the analyzed neurons). We also compared the firing rate before and after HCRT-1 administration with the same approach (Torterolo et al., 2002; Cabrera et al., 2013; Devera et al., 2015; Pascovich et al., 2020). The population mean between these conditions was analyzed using the Wilcoxon Signed-Rank Test. The criterion chosen to discard the null hypothesis was p < 0.05.

## Results

### Recording sites

We recorded 192 neurons that, based on microelectrodes tracks reconstructions, were located within the limits of the mPOA (Figure 1). We excluded seven neurons that were outside this area.

**Figure 1.**
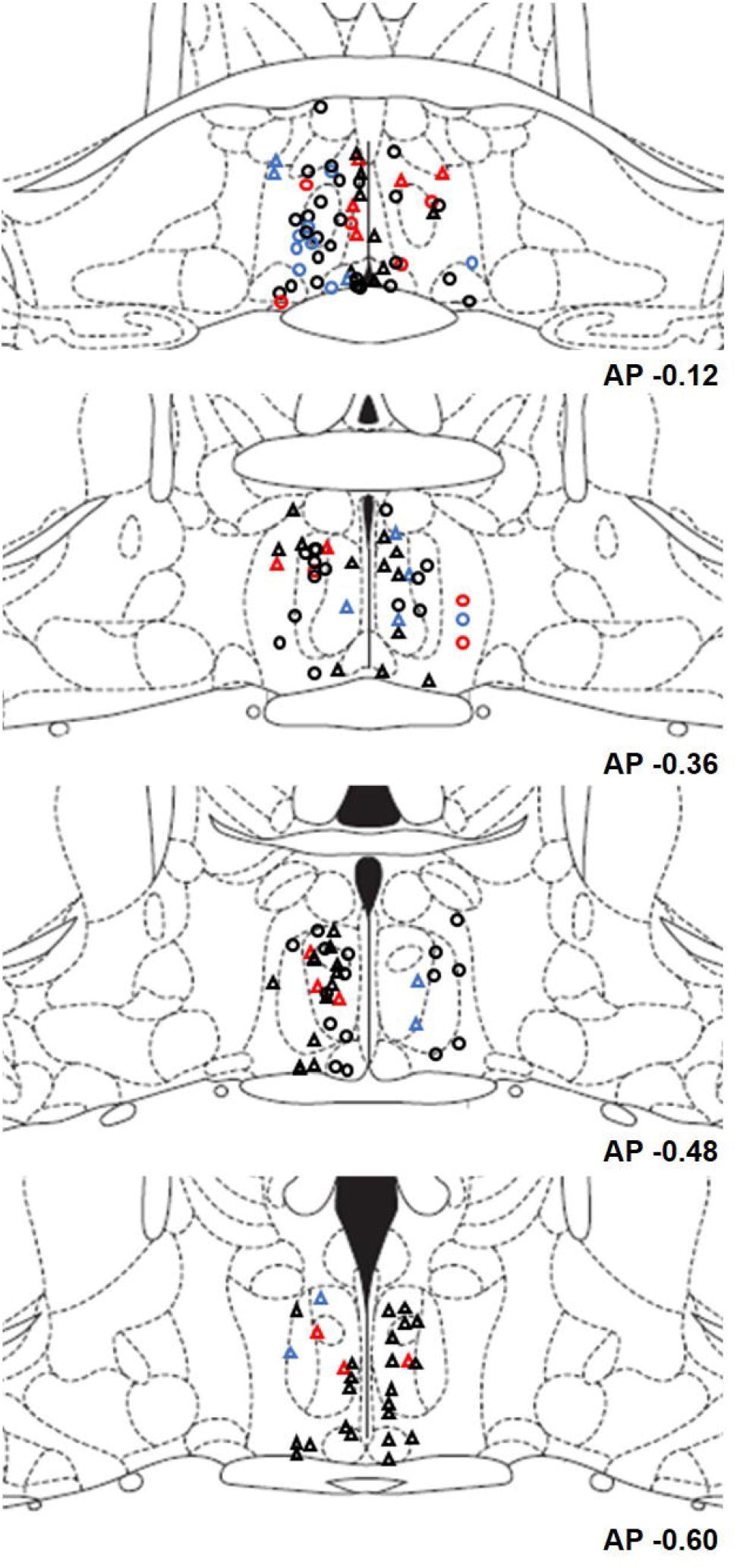
Reconstruction of the recorded neuron sites at the level of mPOA in lactating (circles) and virgin rats (triangles). The colors indicate if neurons increased (red) or decreased (blue) following juxta-cellular administration of HCRT. In black are represented neurons that either did not change their activity following HCRT administration or were not tested. Representative coronal section plates were taken from Paxinos and Watson (2005); the bottom numbers indicate the distance from Bregma.

### Electrophysiological characterization of units in the mPOA

One hundred and four neurons were recorded in 21 lactating rats (4.32 ± 0.14 neurons per animal), and 88 neurons in 15 virgin rats (4.86 ± 0.25 neurons per animal). We first compared the electrophysiological characteristics of the neurons between lactating and virgin rats. We did not find any significant differences in the AP duration, mean and instantaneous frequency, or in the mean or instantaneous frequency CVs (Table 1). However, there was a tendency to increase the AP duration and the instantaneous firing frequency in lactating compared to virgin rats. Lactating and virgin rats also presented equal proportions of neurons with burst firing patterns (68 and 60 %, respectively), defined as two or more spikes with intervals shorter than 10 ms and a decrease in the amplitude of higher-order spikes (Figure 2). The patterns of discharge were also non-statistically different between lactating and virgin rats, with a predominance of the arrhythmic type of discharge (57 % and 61%, respectively) in both conditions.

**Table 1.**
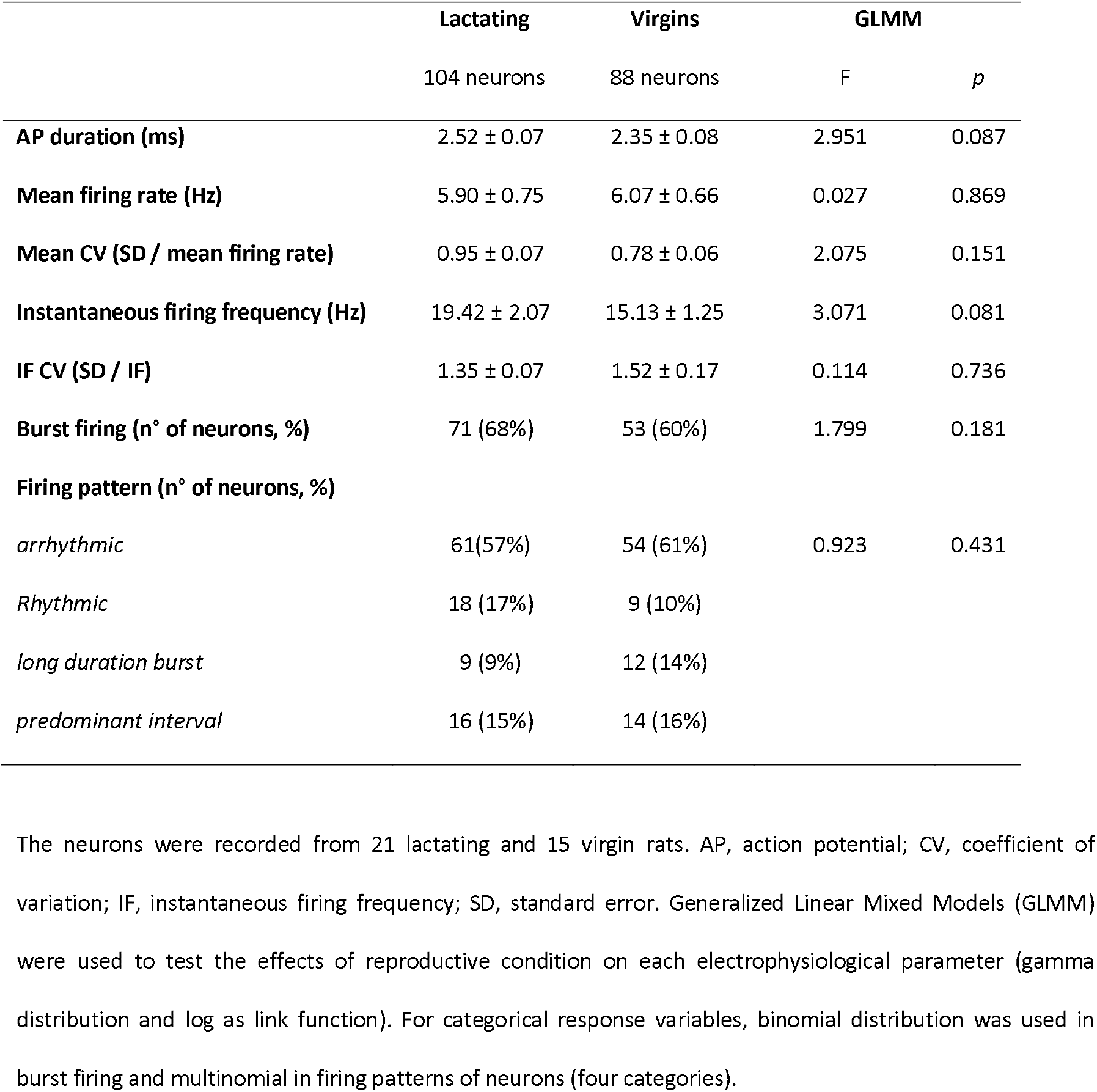
Basic electrophysiological characteristics of mPOA units.

**Figure 2.**
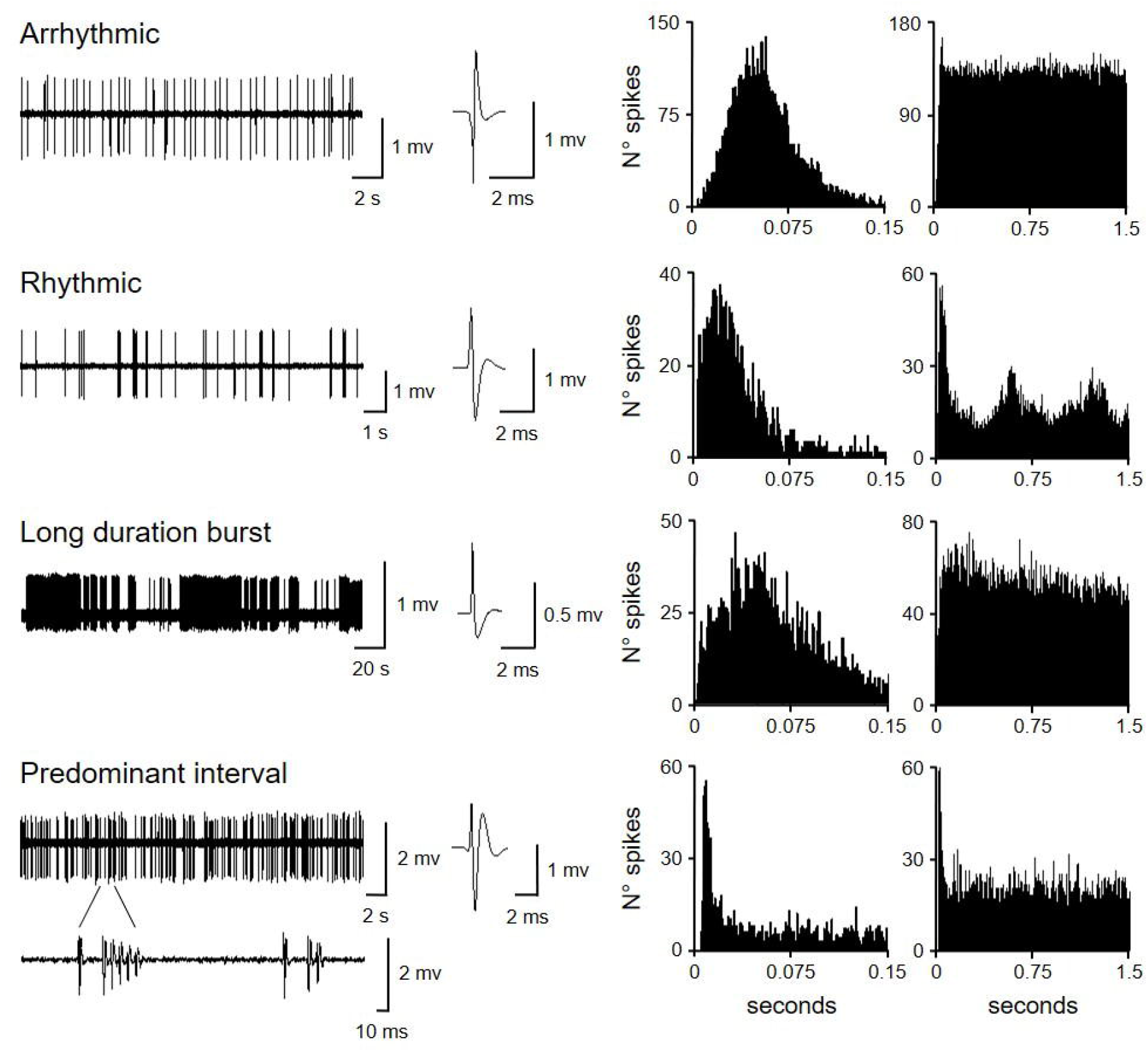
Examples of four neurons with different patterns of discharge. From left to right: raw recording, action potential waveform average, interval histogram, and autocorrelation histogram. The raw recording of the neuron with a predominant interval is shown in a short time frame to highlight the discharge bursts.

### Firing profile of unit recording associated with EEG dynamics

We evaluated 52 mPOA neurons of lactating rats. We found that 29% (n = 15) of these neurons were REMure-ON, 6% (n = 3) were NREMure-ON, while 65% (n = 34) were unrelated to the EEG. In virgin rats, a total of 45 neurons were examined: 13% (n = 6) were REMure-ON, 9% (n = 4) were NREMure-ON and 78% (n = 35) were unrelated to the EEG. This proportion of neurons was not different between lactating and virgin (F = 0.423, p = 0.862). Figure 3 shows examples of each neuronal type.

**Figure 3.**
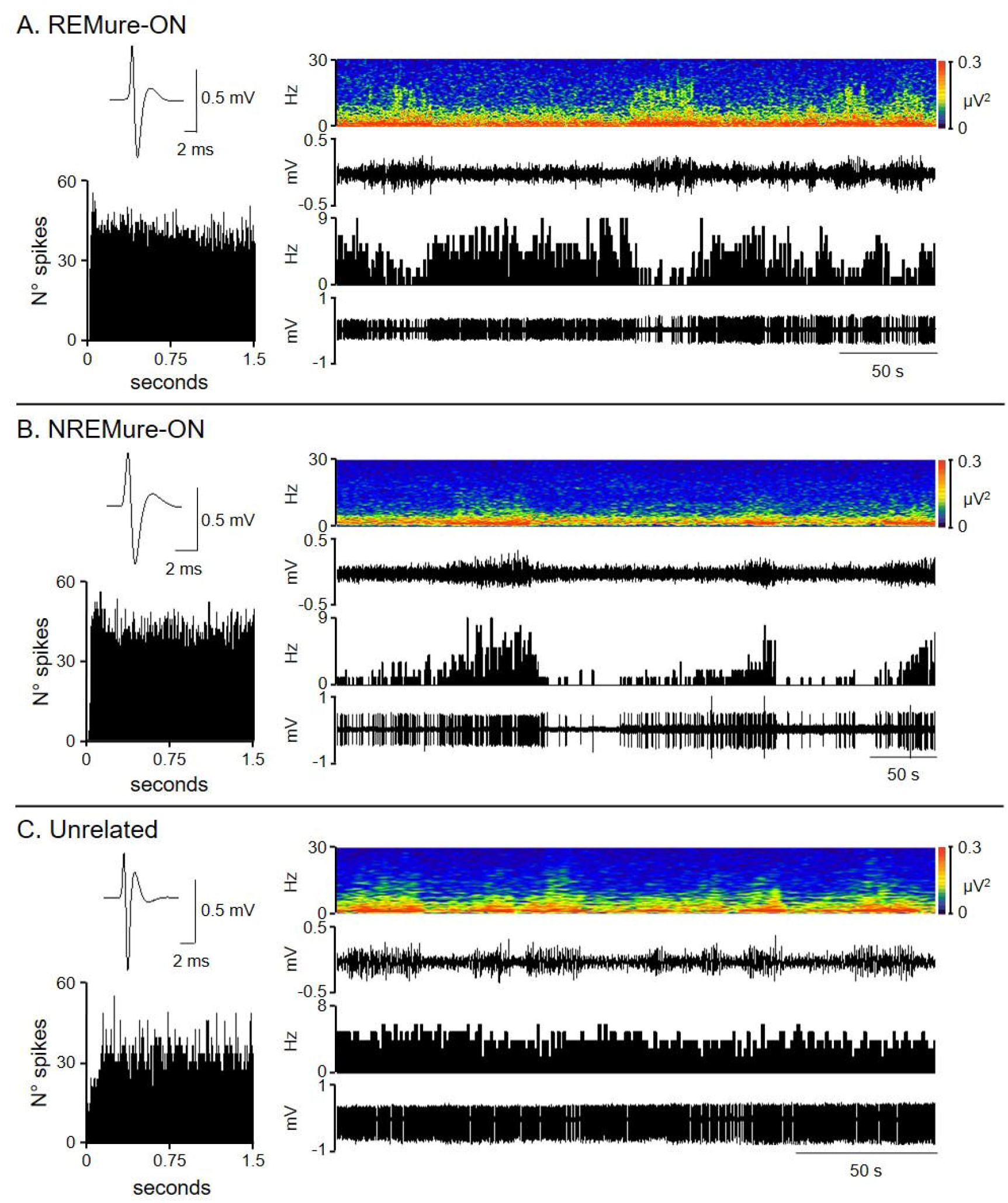
Relationship between mPOA neuronal activity and EEG state. **A**. REMure-ON neuron. The neuron increases its firing rate during EEG activation. **B**. NREMure-ON neuron. This unit increases its activity during EEG with slow waves. **C**. Neuron unrelated with the EEG state. On the left, the waveform average and autocorrelation histogram are shown for each neuron. On the right, from bottom to top: raw unit recording, firing rate histogram, raw EEG, and EEG spectrogram.

The basal mean firing rate of neurons related to the EEG varied significantly among groups (GLMM, F = 3.845 p = 0.025). REMure-ON neurons had lower frequencies compared to those of neurons unrelated to the EEG (t = 3.614, p = 0.012), but not compared to NREMure-ON (t = 5.265, p = 0.302; see Figure 4). We found no differences according to the reproductive condition of animals in the GLMM (F = 0.106, p = 0.746). In addition, AP duration did not vary neither when comparing among the different neuronal types (F = 0.224, p = 0.800), nor when comparing the reproductive condition (F = 3.253, p = 0.074) (Figure 4).

**Figure 4.**
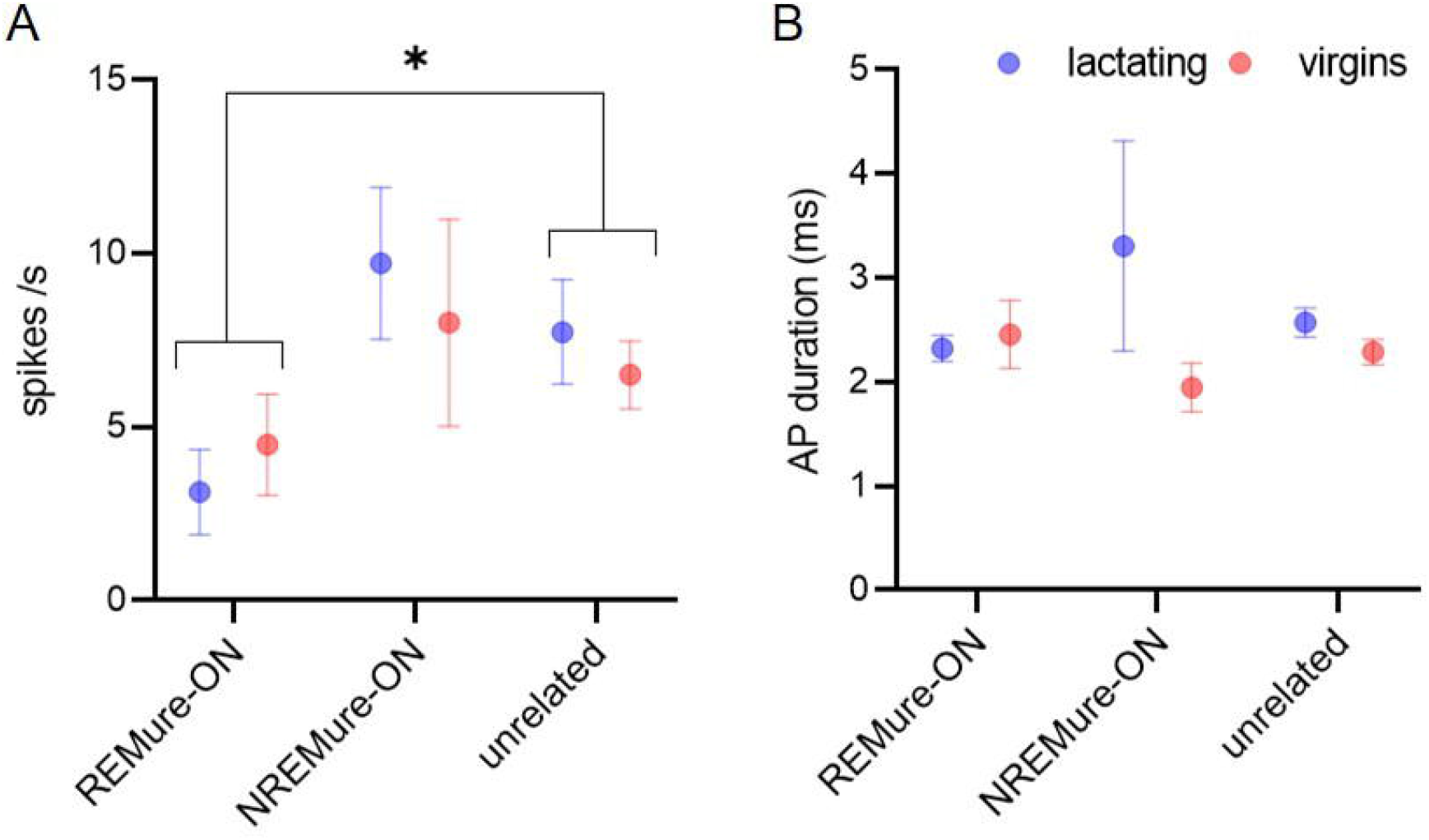
Mean firing rate (A) and action potential duration (B) of neurons according to their relationship with EEG state and reproductive state. * Indicates p < 0.05.

### Effect of HCRT-1

Figure 5 shows examples of the effects of HCRT-1 on the firing rate of mPOA neurons. Juxta-cellular pressure injection of HCRT-1 in lactating rats was achieved in 40 neurons of 16 animals. HCRT-1 increased the firing rate in 13 neurons, from 3.28 ± 0.93 Hz to 5.48 ± 1.46 Hz (W = 91.0, p < 0.001), with a latency of 57.6 ± 16.8 sec and effect duration of 309.3 ± 79.5 sec. HCRT-1 also decreased the firing rate in 13 neurons, from 3.58 ± 1.06 Hz to 1.95 ± 0.78 Hz (W = 91.0, p < 0.001), with a latency of 62.9 ± 11.7 sec and effect duration of 551.0 ± 107.4 sec. Furthermore, HCRT-1 had no effect on the firing rate in 14 neurons (from 10.23 ± 2.42 Hz to 10.14 ± 2.38 Hz; W = 11.0, p = 0.761).

**Figure 5.**
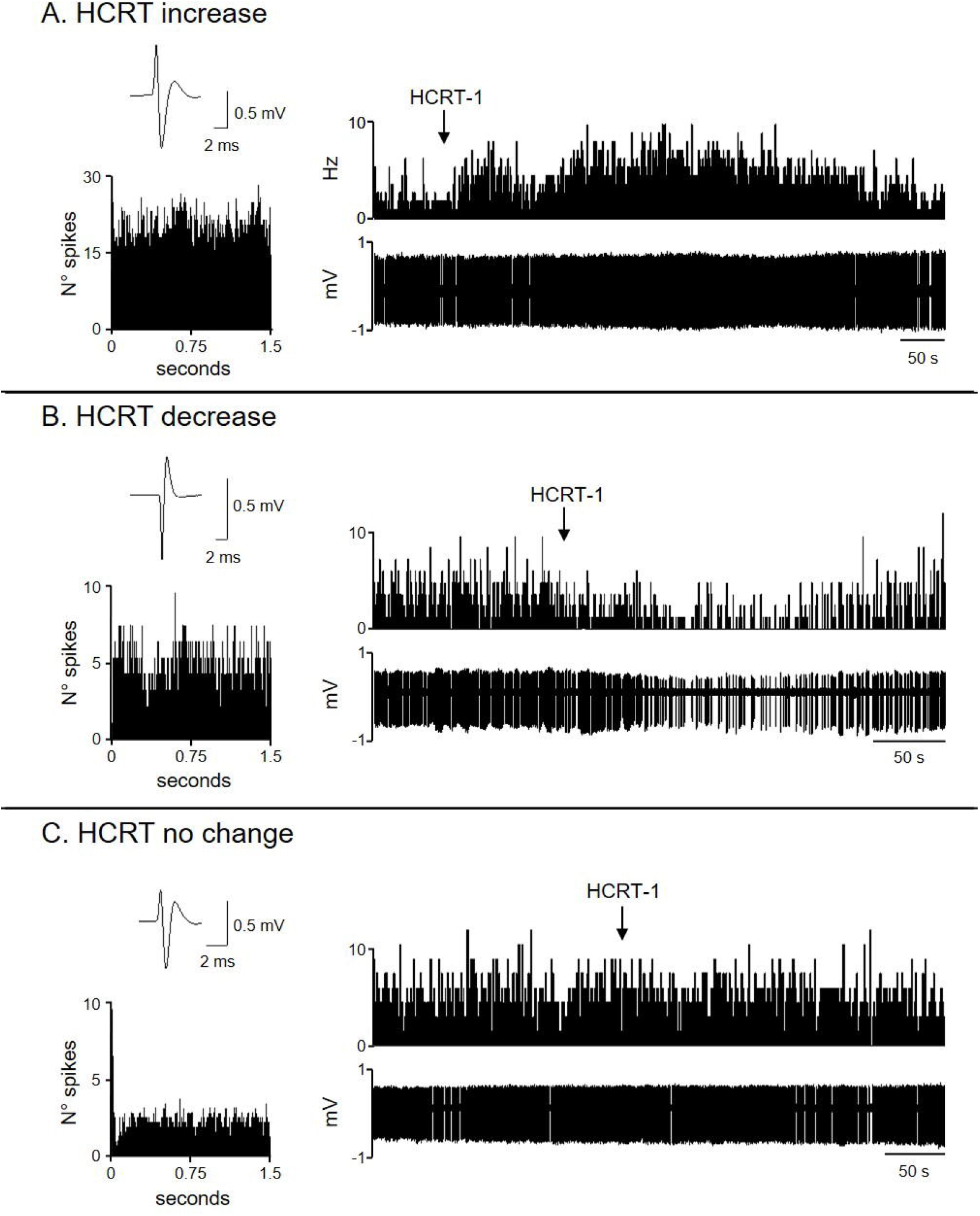
Effects of juxta-cellular administration of HCRT-1 on the activity of mPOA neurons. The raw recording (bottom) and the frequency histogram (top) of three representative neurons were HCRT either increase (A), decrease (B) or had not changed (C) in the neuronal discharge. The arrow shows the time of microinjection of HCRT-1. On the left, the action potential waveform average and autocorrelation histogram for each neuron are shown.

In virgin rats, we injected HCRT-1 in 34 neurons of 14 animals and found that HCRT-1 increased the firing rate in 14 neurons, from 4.09 ± 1.06 Hz to 7.68 ± 2.40 Hz (W = 105.0, p < 0.001), with a latency of 27.4 ± 13.6 sec and effect duration of 188.9 ± 57.7 sec. HCRT-1 also decreased the firing rate in 12 neurons, from 5.53 ± 1.46 Hz to 2.81 ± 0.98 Hz (W = 78.0, p < 0.001) with a latency of 74.6 ± 18.1 sec and effect duration of 307 ± 79.5 sec. Finally, HCRT-1 had no effect in 8 neurons (from 11.18 ± 3.11 to 10.66 ± 2.97; W = 24, p = 0.109).

The proportion of neurons that increased, decreased, or did not change their firing rate following HCRT-1 administration was not different between lactating (32, 32, and 36%, respectively) and virgin rats (39, 33, and 28%, respectively; F = 0.282, p = 0.944).

As a control of the procedure, we analyzed the juxta-cellular administration of saline solution in five neurons of lactating rats, where any modification in their firing rate was seen (from 8.83 ± 1.45 Hz before to 9.25 ± 1.41 Hz after saline administration; T = 0.674, p = 0.500).

The basal mean firing rate of neurons that responded to HCRT-1 juxta-cellular administration was different from those that did not respond, independently of the reproductive condition of animals (GLMM: HCRT effect, F = 10.032, p < 0.001; reproductive condition, F = 1.318, p = 0.255). The basal mean firing rate of neurons that increase (t = 4.273, p < 0.001) or decrease (t = 3.392, p < 0.001) their firing rate following HCRT-1 administration was lower than that of neurons that showed no response to HCRT-1 (Figure 6).

**Figure 6.**
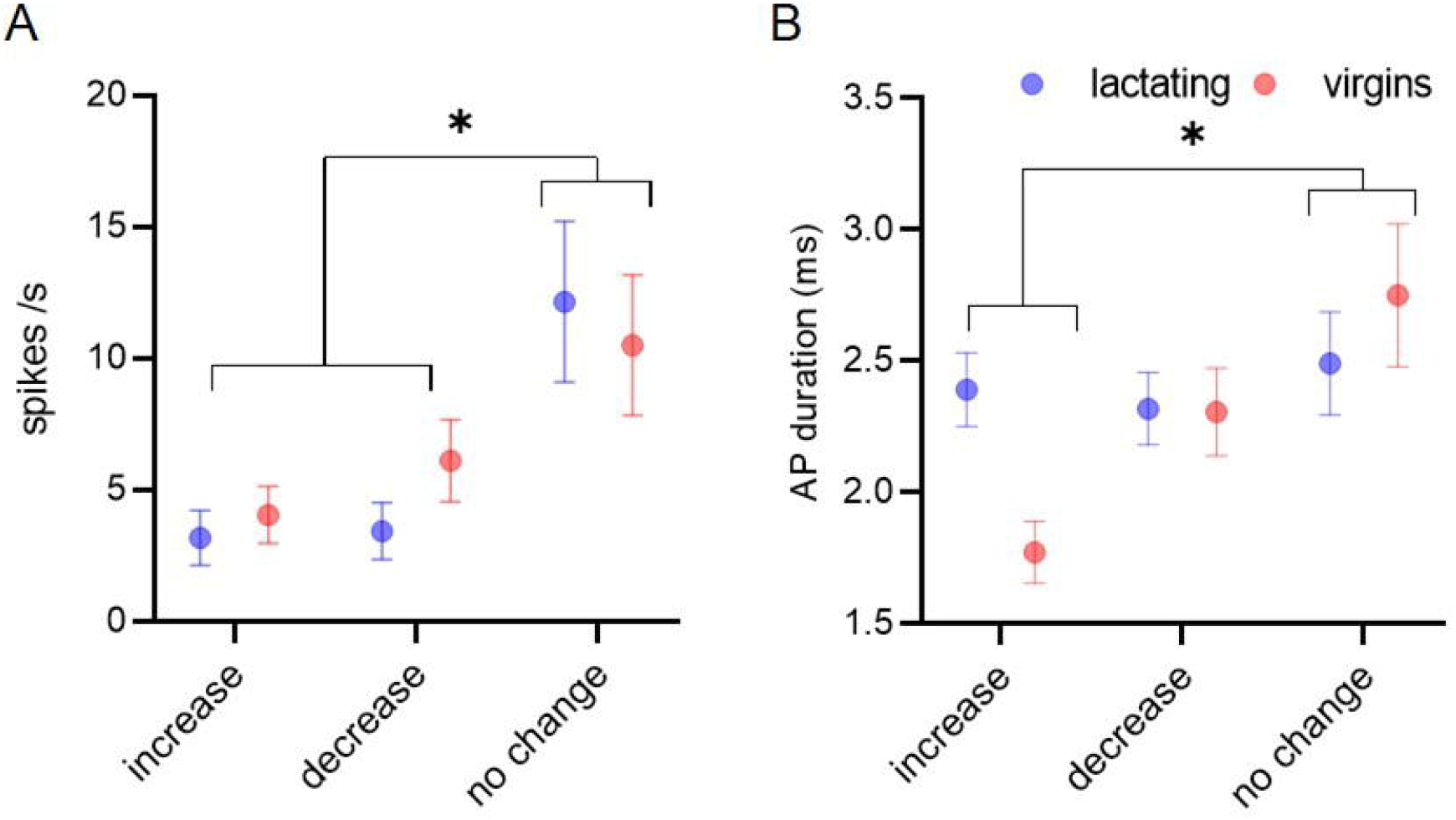
Basal firing rate (A) and action potential duration (B) of neurons according to the response to HCRT-1. Neurons were categorized either as increase, decrease or not change in their mean firing rate, according to the effect of HCRT-1. * Indicates p < 0.05.

The width of the AP had significant variations in the different groups of neurons that responded or not to HCRT-1, being the reproductive condition a non-significant factor in the GLMM (HCRT effect: F = 4.052, p = 0.021; reproductive condition: F = 2.787, p = 0.099). The neurons that responded to HCRT-1 by increasing their firing rate had shorter AP duration than those that did not respond to HCRT-1 (t = -2.803, p = 0.007), without showing significant differences compared to those that decreased their activity (t = 0.273, p = 0.163) (Figure 6).

Independently of reproductive condition of the neurons that increased their firing rate following the microinjection of HCRT-1, 30% were REMure-ON while 0 % were NREMure-ON. From the total of the neurons that decreased their activity in response to HCRT-1, 8 % were REMure-ON while 15 % were NREMure-ON.

## Discussion

In the present study, through *in vivo* extracellular single-unit recordings under urethane anesthesia, we analyzed the electrophysiological characteristics of mPOA neurons in lactating and virgin rats. Moreover, we examined the relationship between the unit activity with the EEG, and the effects of juxta-cellular administration of HCRT-1.

### Characteristics of mPOA neurons in lactating and virgin rats

We found no differences between the main electrophysiological characteristics of mPOA neurons between lactating and virgin rats. Although the duration of AP tended to be longer and the mean instantaneous frequency higher in lactating compared with virgin rats, these differences were not significant, probably caused by the great variability in the recorded neurons.

The electrophysiological properties of mPOA neurons have received little attention during the postpartum period. To our knowledge, no other study has recorded the activity of mPOA neurons in lactating rats. In mice, (Fang et al., 2018) reported that spontaneous firing rate of mPOA neurons was lower in lactating compared to virgin females. This difference between studies may be due to different species or methodological procedures since Fang et al. recordings were done in freely moving conditions. In fact, the mean firing rate was considerably lower (3.36 ± 0.46 and 1.89 ± 0.34 spikes/s, in virgin and lactating rats, respectively) than those observed in the present study (6.07 ± 0.66 and 5.90 ± 0.75 spikes/s, virgin and lactating rats, respectively). In addition, Horrell et al. (2019) using *in vitro* recordings of mPOA neurons from mice males that provide extensive paternal care to their offspring, evidenced that several intrinsic electrophysiological parameters were similar between virgin and father animals.

In agreement with anatomical and neurochemical data (Simerly et al., 1986; Tsuneoka et al., 2013), our data show that mPOA neurons are an electrophysiological heterogeneous group. While we classified most neurons as arrhythmic, there were neurons grouped as “long duration bursts” with rhythmic firing trains separated by long silence intervals; these neurons are consistent with the firing pattern of gonadotropin-releasing hormone (GnRH) neurons of the preoptic area (Suter et al., 2000; Kuehl-Kovarik et al., 2002), suggesting a possible cellular phenotype for these neurons.

### Relationship with the EEG activity

Similarly to our findings, different types of sleep-active neurons (such as NREM-on, REM-on, and NREM/REM-on) has been previously described to coexist within the mPOA (Koyama and Hayaishi, 1994; Suntsova and Dergacheva, 2004; Takahashi et al., 2009). In this sense, in male rats, 14% of recorded neurons were NREM-on, 26% REM-on, and 37% NREM/REM-on (Koyama and Hayaishi, 1994).

For many years, urethane was considered a general anesthetic that resembles the characteristics of physiologic sleep and was utilized as a pharmacological model for its study (Clement et al., 2008; Pagliardini et al., 2013). Hence, as a proxy for the sleep-waking cycle, we utilized urethane anesthesia. We found a higher proportion of REMure-ON neurons (29% and 13%) than NREMure-ON neurons (6% and 4%) in both lactating and virgin rats, respectively. Interestingly, a large percentage of neurons are reported to be active during REM sleep not only in rats (Koyama and Hayaishi, 1994) but also in mice (Takahashi et al., 2009) and cats (Suntsova and Dergacheva, 2004).

Our data exhibit certain differences compared to previous mPOA recordings under urethane anesthesia. Kumar et al. (1989) reported in male rats 13% NREMure-ON and 14 % REMure-ON neurons, while (Lincoln, 1969) in preoptic and anterior hypothalamus of cycling female rats described 27% NREMure-ON and 23% REMure-ON neurons. Besides, Bueno and Pfaff (1976) reported no association between the EEG and the activity of mPOA neurons in ovariectomized rats treated or not with estrogens. Gender, reproductive state, or slight differences in the recording site may account for the differences in the results.

Mondino et al. (2022) have recently described differences between NREMure and REMure states and the corresponding natural sleep states, through a detailed analysis of the EEG that included power, coherence, directionality, and complexity. Given these recent findings, it would be particularly relevant to carry out this characterization of mPOA neurons and the effect of HCRT in non-anesthetized animals.

### HCRT-1 effect in mPOA neurons

Our data show that juxta-cellular HCRT-1 either increases or decreases the firing rate of different mPOA neurons. The proportion of these two groups was similar in lactating and virgin rats. Regarding other subareas of the preoptic area, it has previously been reported that HCRT-1 increases neuronal excitability in median preoptic nucleus (MnPO) neurons, either directly through membrane depolarization via postsynaptic receptor or indirectly through the activation of glutamatergic afferents (Kolaj et al., 2008). As no direct inhibitory effects of HCRT-1 have been reported, we hypothesized that the decrease in the firing rate observed in some neurons was due to presynaptic inhibition through the activation of GABAergic terminals. In this sense, HCRT neurons inhibit the firing of MCH and serotonergic neurons indirectly, increasing the activity of GABAergic neurons (van den Pol et al., 1998; Liu et al., 2002; Apergis-Schoute et al., 2015). Therefore, as previously reported in other brain areas, HCRT _effect_ on mPOA neurons could be mediated either by presynaptic or postsynaptic receptors (van den Pol et al., 1998).

Neurons that HCRT-1 increased their activity had lower basal firing frequencies and narrow AP, compared to those that did not respond to HCRT-1. This fact suggests that HCRT-1 could be acting in a specific group of neurons. However, neurons in which HCRT-1 increased or decreased their activity did not differ from each other, indicating similarities in their electrophysiological characteristics. Given that most preoptic neurons are GABAergic (Lonstein and De Vries, 2000; Tsuneoka et al., 2013; Dimen et al., 2021), this phenotype would be the most likely candidate for most of the recorded neurons. Intracellular *in vitro* recordings within the mPOA show that the firing rate of GABAergic neurons (7.5 ± 3.1 Hz) was greater than the non-GABAergic (3.1 ± 2.9 Hz) (Lundius et al., 2010). However, in the present report, the average firing rate of the neurons that responded to HCRT-1 was 3.43 Hz, similar to the reported for non-GABAergic by (Lundius et al., 2010), suggesting that these neurons are more likely to be non-GABAergic. In this regard, a group of glutamatergic neurons whose activation increases wakefulness and decreases both NREM and REM sleep has been described in the mPOA and surroundings areas (Vanini et al., 2020; Mondino et al., 2021). Given that in previous studies we showed that HCRT-1 within the mPOA of lactating rats increases wakefulness and decreases sleep (Rivas et al., 2021), neurons that increase their activity in response to HCRT-1 may be glutamatergic. However, a phenotypic identification is necessary to confirm this hypothesis.

In neurons where HCRT increases its firing rate, 30% were REMure-ON while none was NREMure-ON. Given the wake-promoting role of HCRT, we might expect that HCRT activates neurons that are wake-ON. Since these recordings were made under anesthesia, we cannot determine if the REMure-ON neurons were either wake-ON or REM-ON, although Mondino et al. (2022) stated that the active state under urethane has more similarities to REM sleep than to wakefulness. On the other hand, from the total of neurons that HCRT decreased its activity, 8% were REMure-ON while 15% were NREMure-ON. The fact that HCRT decreased the activity of NREMure-ON neurons (presumably active NREM neurons) is consistent with the wake-promoting role of HCRT and with the reciprocal inhibition between sleep and wake-promoting systems that has been suggested (Saper et al., 2010). In the latter sense, reciprocal anatomical and functional connections between VLPO sleep-promoting galaninergic inhibitory neurons and monoaminergic, cholinergic, and lateral hypothalamic wake-promoting neurons have been described (Gallopin et al., 2000; Venner et al., 2019). Interestingly, HCRTergic neurons are also inhibited by sleep-active preoptic neurons (Saito et al., 2013).

Our previous evidence shows that HCRT in the mPOA promote wakefulness, along with some components of maternal behavior, and increase body temperature (Rivas et al., 2016; Rivas et al., 2021). However, the specific neuronal networks that mediate these physiological effects, and the type of mPOA neurons involved remain unknown. Since the neurochemical phenotype of mPOA neurons includes a great variety of possible neurotransmitters (Simerly et al., 1986; Tsuneoka et al., 2013), the future identification of recorded neurons in which HCRT-1 modulates its activity is fundamental to understand the functions of the hypocretinergic projections towards the mPOA.

## Conclusions

The results of our extracellular *in vivo* recordings indicate that the electrophysiological characteristics of mPOA neurons did not differ between postpartum and virgin rats. Moreover, some of these neurons showed an EEG-related activity, in similar proportions between virgin and lactating rats. In addition, the juxta-cellular HCRT administration increased the activity of one group of neurons while decreasing it in others, both in postpartum and virgin rats. Finally, these HCRT-1-responsive neurons exhibited distinct electrophysiological characteristics from those that did not respond.

This is the first study that shows an electrophysiological characterization of mPOA neurons as well as the effect of HCRT on these neurons in lactating rats. Since the effects of HCRT on mPOA neurons were not the same in all neurons, this study suggests that the mechanisms by which HCRT modulate the different parameters and behaviors controlled by the mPOA include different cell population.

## Acknowledgements

This work was partially supported by ‘‘Programa de Desarrollo de Ciencias Básicas (PEDECIBA), UdelaR” and ‘‘Agencia Nacional de Investigación e Innovación (ANII)”.

## References

Agresti, A., 2003. Categorical data analysis, Vol., John Wiley & Sons.

Akbari, E.M., et al., 2013. The effects of parity and maternal behavior on gene expression in the medial preoptic area and the medial amygdala in postpartum and virgin female rats: A microarray study. Behav Neurosci. 127, 913–22.

Apergis-Schoute, J., et al., 2015. Optogenetic evidence for inhibitory signaling from orexin to MCH neurons via local microcircuits. J Neurosci. 35, 5435–41.

Benedetto, L., et al., 2021. Local administration of bicuculline into the ventrolateral and medial preoptic nuclei modifies sleep and maternal behavior in lactating rats. Physiology & Behavior. 238, 113491.

Bosch, O.J., Neumann, I.D., 2008. Brain vasopressin is an important regulator of maternal behavior independent of dams’ trait anxiety. Proc Natl Acad Sci U S A. 105, 17139–44.

Boutrel, B., Cannella, N., de Lecea, L., 2010. The role of hypocretin in driving arousal and goal-oriented behaviors. Brain Res. 1314, 103–11.

Bueno, J., Pfaff, D.W., 1976. Single unit recording in hypothalamus and preoptic area of estrogen-treated and untreated ovariectomized female rats. Brain Res. 101, 67–78.

Cabrera, G., et al., 2013. Wakefulness-promoting role of the inferior colliculus. Behav Brain Res. 256, 82–94.

Clement, E.A., et al., 2008. Cyclic and sleep-like spontaneous alternations of brain state under urethane anaesthesia. PLoS One. 3, e2004.

D’Anna, K.L., Gammie, S.C., 2006. Hypocretin-1 dose-dependently modulates maternal behaviour in mice. J Neuroendocrinol. 18, 553–66.

Devera, A., et al., 2015. Melanin-concentrating hormone (MCH) modulates the activity of dorsal raphe neurons. Brain Res. 1598, 114–28.

Dimen, D., et al., 2021. Sex-specific parenting and depression evoked by preoptic inhibitory neurons. iScience. 24, 103090.

Dobolyi, A., 2009. Central amylin expression and its induction in rat dams. J Neurochem. 111, 1490–500.

Driessen, T.M., et al., 2014. Endogenous CNS expression of neurotensin and neurotensin receptors is altered during the postpartum period in outbred mice. PLoS One. 9, e83098.

Espana, R.A., Berridge, C.W., Gammie, S.C., 2004. Diurnal levels of Fos immunoreactivity are elevated within hypocretin neurons in lactating mice. Peptides. 25, 1927–34.

Fang, Y.Y., et al., 2018. A Hypothalamic Midbrain Pathway Essential for Driving Maternal Behaviors. Neuron. 98, 192–207 e10.

Fleming, A.S., Walsh, C., 1994. Neuropsychology of maternal behavior in the rat: c-fos expression during mother-litter interactions. Psychoneuroendocrinology. 19, 429–43.

Gallopin, T., et al., 2000. Identification of sleep-promoting neurons in vitro. Nature. 404, 992–5.

Gammie, S.C., 2005. Current models and future directions for understanding the neural circuitries of maternal behaviors in rodents. Behav Cogn Neurosci Rev. 4, 119–35.

Gong, H., et al., 2004. Activation of c-fos in GABAergic neurones in the preoptic area during sleep and in response to sleep deprivation. J Physiol. 556, 935–46.

Hajos, M., et al., 1995. Evidence for a repetitive (burst) firing pattern in a sub-population of 5-hydroxytryptamine neurons in the dorsal and median raphe nuclei of the rat. Neuroscience. 69, 189–97.

Harris, G.C., Wimmer, M., Aston-Jones, G., 2005. A role for lateral hypothalamic orexin neurons in reward seeking. Nature. 437, 556–9.

Horrell, N.D., Saltzman, W., Hickmott, P.W., 2019. Plasticity of paternity: Effects of fatherhood on synaptic, intrinsic and morphological characteristics of neurons in the medial preoptic area of male California mice. Behav Brain Res. 365, 89–102.

Insel, T.R., 1990. Regional changes in brain oxytocin receptors post-partum: time-course and relationship to maternal behaviour. J Neuroendocrinol. 2, 539–45.

Kolaj, M., Coderre, E., Renaud, L.P., 2008. Orexin peptides enhance median preoptic nucleus neuronal excitability via postsynaptic membrane depolarization and enhancement of glutamatergic afferents. Neuroscience. 155, 1212–20.

Koyama, Y., Hayaishi, O., 1994. Firing of neurons in the preoptic/anterior hypothalamic areas in rat: its possible involvement in slow wave sleep and paradoxical sleep. Neurosci Res. 19, 31–8.

Kuehl-Kovarik, M.C., et al., 2002. Episodic bursting activity and response to excitatory amino acids in acutely dissociated gonadotropin-releasing hormone neurons genetically targeted with green fluorescent protein. J Neurosci. 22, 2313–22.

Kumar, V.M., Datta, S., Singh, B., 1989. The role of reticular activating system in altering medial preoptic neuronal activity in anesthetized rats. Brain Res Bull. 22, 1031–7.

Kumar, V.M., 2004. Why the medial preoptic area is important for sleep regulation. Indian J Physiol Pharmacol. 48, 137–49.

Lincoln, D.W., 1969. Correlation of unit activity in the hypothalamus with EEG patterns associated with the sleep cycle. Exp Neurol. 24, 1–18.

Liu, R.J., van den Pol, A.N., Aghajanian, G.K., 2002. Hypocretins (orexins) regulate serotonin neurons in the dorsal raphe nucleus by excitatory direct and inhibitory indirect actions. J Neurosci. 22, 9453–64.

Lonstein, J.S., et al., 1998. Forebrain expression of c-fos due to active maternal behaviour in lactating rats. Neuroscience. 82, 267–81.

Lonstein, J.S., De Vries, G.J., 2000. Maternal behaviour in lactating rats stimulates c-fos in glutamate decarboxylase-synthesizing neurons of the medial preoptic area, ventral bed nucleus of the stria terminalis, and ventrocaudal periaqueductal gray. Neuroscience. 100, 557–68.

Lundius, E.G., et al., 2010. Histamine influences body temperature by acting at H1 and H3 receptors on distinct populations of preoptic neurons. J Neurosci. 30, 4369–81.

Mann, P.E., Bridges, R.S., 2001. Lactogenic hormone regulation of maternal behavior. Prog Brain Res. 133, 251–62.

McGregor, R., et al., 2011. Highly specific role of hypocretin (orexin) neurons: differential activation as a function of diurnal phase, operant reinforcement versus operant avoidance and light level. J Neurosci. 31, 15455–67.

Mondino, A., et al., 2021. Glutamatergic neurons in the preoptic hypothalamus promote wakefulness, destabilize NREM sleep, suppress REM sleep, and regulate cortical dynamics. J Neurosci.

Mondino, A., et al., 2022. Urethane anaesthesia exhibits neurophysiological correlates of unconsciousness and is distinct from sleep. European Journal of Neuroscience. n/a 1–19.

Muschamp, J.W., et al., 2007. A role for hypocretin (orexin) in male sexual behavior. J Neurosci. 27, 2837–45.

Numan, M., 1974. Medial preoptic area and maternal behavior in the female rat. J Comp Physiol Psychol. 87, 746–59.

Numan, M., et al., 1999. Expression of intracellular progesterone receptors in rat brain during different reproductive states, and involvement in maternal behavior. Brain Res. 830, 358–71.

Numan, M., 2006. Hypothalamic neural circuits regulating maternal responsiveness toward infants. Behav Cogn Neurosci Rev. 5, 163–90.

Pagliardini, S., Gosgnach, S., Dickson, C.T., 2013. Spontaneous sleep-like brain state alternations and breathing characteristics in urethane anesthetized mice. PLoS One. 8, e70411.

Parent, C., et al., 2017. Maternal care associates with differences in morphological complexity in the medial preoptic area. Behav Brain Res. 326, 22–32.

Pascovich, C., et al., 2020. Melanin-concentrating hormone (MCH) in the median raphe nucleus: Fibers, receptors and cellular effects. Peptides. 126, 170249.

Paxinos, G., Watson, C., 2005. The Rat Brain in Stereotaxic Coordinates, Vol., Elsevier Academic Press, San Diego, California.

Peyron, C., et al., 1998. Forebrain afferents to the rat dorsal raphe nucleus demonstrated by retrograde and anterograde tracing methods. Neuroscience. 82, 443–68.

Rivas, M., et al., 2016. Hypocretinergic system in the medial preoptic area promotes maternal behavior in lactating rats. Peptides. 81, 9–14.

Rivas, M., et al., 2021. Role of Hypocretin in the Medial Preoptic Area in the Regulation of Sleep, Maternal Behavior and Body Temperature of Lactating Rats. Neuroscience. 475, 148–162.

Rondini, T.A., et al., 2010. Chemical identity and connections of medial preoptic area neurons expressing melanin-concentrating hormone during lactation. J Chem Neuroanat. 39, 51–62.

Saito, Y.C., et al., 2013. GABAergic neurons in the preoptic area send direct inhibitory projections to orexin neurons. Front Neural Circuits. 7, 192.

Saper, C.B., et al., 2010. Sleep state switching. Neuron. 68, 1023–42.

Shams, S., et al., 2012. Dendritic morphology in the striatum and hypothalamus differentially exhibits experience-dependent changes in response to maternal care and early social isolation. Behav Brain Res. 233, 79–89.

Simerly, R.B., Gorski, R.A., Swanson, L.W., 1986. Neurotransmitter specificity of cells and fibers in the medial preoptic nucleus: an immunohistochemical study in the rat. J Comp Neurol. 246, 343–63.

Stack, E.C., Numan, M., 2000. The temporal course of expression of c-Fos and Fos B within the medial preoptic area and other brain regions of postpartum female rats during prolonged mother--young interactions. Behav Neurosci. 114, 609–22.

Suntsova, N.V., Dergacheva, O.Y., 2004. The role of the medial preoptic area of the hypothalamus in organizing the paradoxical phase of sleep. Neurosci Behav Physiol. 34, 29–35.

Suter, K.J., et al., 2000. Whole-cell recordings from preoptic/hypothalamic slices reveal burst firing in gonadotropin-releasing hormone neurons identified with green fluorescent protein in transgenic mice. Endocrinology. 141, 3731–6.

Szymusiak, R., et al., 1998. Sleep-waking discharge patterns of ventrolateral preoptic/anterior hypothalamic neurons in rats. Brain Res. 803, 178–88.

Takahashi, K., Lin, J.S., Sakai, K., 2009. Characterization and mapping of sleep-waking specific neurons in the basal forebrain and preoptic hypothalamus in mice. Neuroscience. 161, 269–92.

Torterolo, P., et al., 1998. Auditory cortical efferent actions upon inferior colliculus unitary activity in the guinea pig. Neurosci Lett. 249, 172–6.

Torterolo, P., et al., 2002. Inferior colliculus unitary activity in wakefulness, sleep and under barbiturates. Brain Res. 935, 9–15.

Torterolo, P., et al., 2003. Hypocretinergic neurons are primarily involved in activation of the somatomotor system. Sleep. 26, 25–8.

Torterolo, P., et al., 2011. Hypocretinergic neurons are activated in conjunction with goal-oriented survival-related motor behaviors. Physiol Behav. 104, 823–30.

Trivedi, P., et al., 1998. Distribution of orexin receptor mRNA in the rat brain. FEBS Lett. 438, 71–5.

Tsuneoka, Y., et al., 2013. Functional, anatomical, and neurochemical differentiation of medial preoptic area subregions in relation to maternal behavior in the mouse. J Comp Neurol. 521, 1633–63.

van den Pol, A.N., et al., 1998. Presynaptic and postsynaptic actions and modulation of neuroendocrine neurons by a new hypothalamic peptide, hypocretin/orexin. J Neurosci. 18, 7962–71.

Vanini, G., et al., 2020. Activation of Preoptic GABAergic or Glutamatergic Neurons Modulates Sleep-Wake Architecture, but Not Anesthetic State Transitions. Curr Biol. 30, 779–787 e4.

Vanini, G., Torterolo, P., 2021. Sleep-Wake Neurobiology. Adv Exp Med Biol. 1297, 65–82.

Venner, A., et al., 2019. An Inhibitory Lateral Hypothalamic-Preoptic Circuit Mediates Rapid Arousals from Sleep. Curr Biol. 29, 4155–4168 e5.

Wagner, C.K., Morrell, J.I., 1996. Levels of estrogen receptor immunoreactivity are altered in behaviorally-relevant brain regions in female rats during pregnancy. Brain Res Mol Brain Res. 42, 328–36.

Wang, J.B., et al., 2003. Variation in the expression of orexin and orexin receptors in the rat hypothalamus during the estrous cycle, pregnancy, parturition, and lactation. Endocrine. 22, 127–34.

